# DIS3L2 is essential for neural crest survival by modulating Akt signaling

**DOI:** 10.1101/2024.12.26.630456

**Authors:** Sian D’Silva, Tuhina Prasad, Megha Kumar

**Affiliations:** CSIR-Centre for Cellular and Molecular Biology (CSIR-CCMB), Habsiguda, Uppal road, Hyderabad-500007, India; Academy of Scientific and Innovative Research (AcSIR), Ghaziabad-201002, India

**Author notes:** Both authors contributed equally to this work (Sian D’Silva, Tuhina Prasad).

**Keywords:** DIS3L2, apoptosis, neural crest, Akt signaling, zebrafish

## Abstract

DIS3 like 3’-5’ exoribonuclease 2 (DIS3L2), an exoribonuclease is known to preferentially degrade uridylated RNA substrates, miRNAs, and ncRNAs. Recent reports show that DIS3L2 also plays a key role in cell proliferation and tumor growth. Mutations in *DIS3L2* are associated with congenital disorders such as Perlman syndrome, yet the developmental functions of DIS3L2 remain unknown. We report the developmental role of *dis3l2* in neural crest specification, patterning, and survival in the zebrafish embryo. *Dis3l2* morphants exhibited reduced expression of neural crest specifier genes coupled with extensive apoptosis in the neural tissue. Our study demonstrates that DIS3L2 regulates neuronal apoptosis and progenitor functions through the Akt –GSK3β signaling pathway. Additionally, we show that DIS3L2 is essential for early mitoses in the zebrafish blastula and plays a key role in maintaining spindle length at metaphase, chromosome congression, spindle pole integrity, and cytokinesis. In summary, we identify new functions of exoribonuclease DIS3L2 in cell fate specification, neural progenitor survival, and mitosis during embryogenesis which form the underlying basis of *DIS3L2-associated* overgrowth Perlman syndrome.

**Plain English summary:** DIS3 like 3’-5’ exoribonuclease 2 (DIS3L2) degrades mRNAs and various RNA substrates in the eukaryotic cells. DIS3L2 mutations are associated with congenital overgrowth syndromes like Perlman syndrome and Wilm’s tumor. The role of DIS3L2 in regulating the embryonic processes remains poorly understood. Our study delineates the developmental functions of DIS3L2 in cell fate determination, survival, and proliferation during vertebrate embryogenesis using zebrafish embryos as a model. We show that DIS3L2 plays a key role in neural crest survival and patterning by modulating Akt-GSKβ signaling. We also report unique molecular functions of DIS3L2 in mitotic fidelity and cytokinesis during embryonic mitoses. Our study provides novel insights into the molecular functions of DIS3L2 in regulating neuronal apoptosis during embryonic brain morphogenesis and associated CNS disorders.

## Background

DIS3 like 3’-5’ exoribonuclease 2 (DIS3L2), is a member of the RNase II family of 3’-5’ exoribonucleases which binds to the core RNA exosome complex. DI3L2 can degrade both single and double-stranded RNA substrates *in vitro* and is known to preferentially degrade uridylated RNA substrates in addition to degradation of mature miRNAs, aberrant mRNAs, and non-coding RNAs (1). A recent report shows that DIS3L2 can induce apoptotic mRNA decay and inhibition of translation (2). It has many biological functions including cell division and differentiation. *In vitro* studies show that *DIS3L2* knockdown promotes cell proliferation and cancer cell growth by regulating cell cycle progression (3). Particularly, siRNA-mediated knockdown in HeLa cells shows that depletion of *DIS3L2* results in mitotic errors, and multiple spindle poles. *DIS3L2* regulates size and proliferation in the developing imaginal discs in *Drosophila* by upregulating Igdf2 signaling (4–6). The *DIS3L2*-depleted mice show Perlman syndrome-associated features and the homozygous *DIS3L2* null mice failed to survive the first postnatal day due to underinflated lungs, hypotonia, bradykinesia, and abnormal spine curvature (1,3). However, the developmental functions of DIS3L2 remain unknown to date. Our study delineates the functional role of DIS3L2 in cranial neural crest specification, patterning, and survival during early embryogenesis. We demonstrate that DIS3L2 functions via Akt-GSK3β signaling to modulate cell fate specification, apoptosis, and cell survival in the neural crest cells. Further, DIS3L2 is vital for early mitotic events in the zebrafish blastulae, ensuring proper chromosome alignment, spindle pole stability, metaphase spindle length, and cytokinesis.

## Methods

### Zebrafish lines, MO injection, and characterization of phenotypes

Tubingen strain (TU-AB) zebrafish were raised according to standard protocols described earlier (51). All experiments were performed according to protocols approved by the Institutional Animal Ethics Committee of the Council of Scientific and Industrial Research Centre for Cellular and Molecular Biology, India. Embryos were obtained from the natural spawning of adult fish, kept at 28.5°C, and staged according to hours and days after fertilization (52). The endogenous DIS3L2 levels were depleted by microinjecting 0.1mM, 0.5mM or 1mM DIS3L2 translation blocker morpholino (Gene Tools), 5’-ATGCCATAATCTCGTCTACAGTTAC-3′ at the one-cell stage. The embryos were then analyzed for gross morphological defects, and survival at later stages of development (24hpf).

### RNA isolation, Semi□quantitative RT□PCR, and sequencing

Total RNA was isolated from embryos using the RNA isolation kit (MN; 740955.50) as per the manufacturer’s protocol. 1 μg of total RNA was taken to synthesize cDNA by using PrimeScript 1st strand cDNA synthesis kit (Takara, Cat. No. 6110A). RT-PCR was carried out for the multiple developmental stages for amplification of zebrafish DIS3L2 with *beta-actin* as a control. PCR was performed in BioRad C1000 Touch thermal cycler with, Taq DNA Polymerase (TaKaRa, cat. no. R500A). The following PCR conditions were used for amplification of *dis3l2*: initial denaturation was at 94 °C for 5 min, followed by 30 cycles of denaturation at 94 °C for 30 secs, annealing at 60°C for 30 secs, and extension at 72 °C for 30 sec. The final extension was for 10 min at 72 °C. Amplified PCR products were examined and pictures were captured using the Vilber-Lourmat Gel Documentation system (Germany).

### Real-time PCR

For quantitative real-time PCR (qPCR), we performed PCR using Power SYBR Green PCR Master Mix (Applied Biosystems) with ViiA 7 Real-Time PCR System (Applied Biosystems) according to the manufacturer’s instructions. 100 ng/μL cDNA of control and injected embryos were used along with PowerSYBR Green PCR master mix (Applied Biosystems, Cat. No. 4367659), and the following program was used: 40 cycles of 95 °C−10 min, 95 °C−15 s, 60 °C−1 min. Quantification data are represented as means ± SEM of three independent experiments. Analyses on normalized data were performed using the 2−δδCT algorithm (the delta−delta-Ct or ddCt algorithm). All genes were normalized against *beta-actin* unless mentioned otherwise. The GraphPad Prism software was used for the analysis. For the statistical significance of the data, p-values were calculated by performing an unpaired Student t-test. All primers used in this study are listed in Fig.S1 table.

### Antisense riboprobe preparation

The total RNA was used to prepare cDNA using the PrimeScript 1^st^ strand cDNA synthesis kit (TaKaRa, Cat. # 6110A). Sequence-specific primers were used to amplify the specific gene fragment and cloned into the pGEMT easy vector and the sequence was verified. The sequence-verified plasmid was linearised using Nco1 and antisense digoxigenin (DIG)--labeled riboprobes were synthesized using the DIG RNA labeling kit (Roche #11175025910). (51).

### Whole Mount RNA in Situ Hybridization

Control and injected embryos at 1dpf were fixed in 4% paraformaldehyde overnight followed by methanol fixation. Methanol-fixed embryos were rehydrated with 50% MeOH/PBT (1× PBS and 0.1% Tween 20). Embryos were rinsed with PBT, a 1:1 PBT/hybridization wash, and then a hybridization wash (50% formamide, 1.3 × SSC, 5 mM EDTA, 0.2%, Tween-20, H2O). Embryos were incubated with a hybridization mix (50% formamide, 1.3 × SSC, 5 mM EDTA, 0.2%, Tween-20, 50 μg/mL yeast t RNA, 100 μg/mL heparin, H2O) for 1 h at 55 °C. The riboprobes prepared in the hybridization mix were added and incubated for 16 h at 55 °C. Then, embryos were rinsed with prewarmed hybridization wash for 30 min (2 times) at 55 °C and followed by 1 × TBST wash (5 M NaCl, 1 M KCl, 1 M Tris, pH 7.5, and 10% Tween 20) at room temperature. Then, blocking was done with 10% FBS heat-inactivated fetal bovine serum (Gibco; 16210-064) for 1 h at room temperature. The embryos were then incubated with an Anti-DIG-AP antibody (Roche 11093274910) with a dilution of 1:5000 overnight at 4°C. The next day, embryos were transferred into a 6-well plate and washed twice with 1X NTMT (0.1M NaCl, 0.1 M Tris-Cl at pH 9.5, 0.05M MgCl2, 1% Tween-20 and H2O) for 10 min; then, incubated with 1:1 NBT-BCIP solution (Cat no. PI34042) NTMT solution for the color development. The embryos were monitored during this color reaction and stopped with 1× PBT as a stop solution. Images of embryos were captured using a Zeiss Stemi 508 stereomicroscope.

### Western Blot analysis

Zebrafish embryos (n=100) were manually dechorinated and deyolked with a deyolking buffer and gentle pipetting. The deyolked embryos were then washed followed by 100ul of Laemmli buffer and homogenized with a 2ml syringe. The lysates were boiled and then subjected to SDS-PAGE. 20 μg of total protein was loaded in each lane on 8% SDS-PAGE gel for detection of the desired proteins. Proteins were transferred to PVDF membranes (Merck Milipore, ISEQ00010) by electro-blotting. Membranes were blocked for 2 h at room temperature in 5% Blotto non-fat dry milk/TBST (20 mM Tris HCl, pH 7.5, 150 mM NaCl, 0.1% TWEEN 20) and probed overnight with primary antibodies. After three washes with TBST, the blots were incubated with HRP-conjugated secondary antibody for 1 hr, and then washed in TBST. The blots were developed for chemiluminescence signal using the chemiluminescence (Bio-Rad) and captured in the Vilber-Lourmat Chemiluminescence System (Germany). The indicated antibody dilutions were used for western blot. Rabbit antibodies: anti-DIS3L2 (1:2000) (Thermo, #PA5-31273), anti-CASPASE 3 (1:1000) (CST# 9662), anti-CASPASE 9 (1:000) (Thermo # PA5-22252), anti-GSK3β (1:1000) (CST#1256), anti-pGSK3β ser9 (1:1000) (CST#5558). Mouse antibodies: anti-CDK1 (1:1000), (Invitrogen #33-1800), anti-Mek1 (1:1000) (Invitrogen#13-3500), anti-GAPDH (1:2000) (Invitrogen #MA5-15738), anti-β tubulin (1:2000) (Thermo#MA5-16308), anti-AKT (1:1000) (CST#40D4, 2920), anti-pAKT thr308 (1:1000) (CST#D25E6, 13038).

### Immunohistochemistry of zebrafish embryos

Embryos at 256 cell stage were fixed in 4% paraformaldehyde overnight followed by methanol at −20 °C. Methanol-fixed embryos were rehydrated with 50% MeOH/PBST (1X PBS and 0.1% Triton X-100) followed by incubation in blocking solution 1% BSA/PBST for 2 h at room temperature. Then the embryos were incubated with primary antibody overnight at 4 °C. The following day embryos were washed with 1X PBST and then incubated overnight at 4 °C with the appropriate secondary antibody. The next day embryos are washed with 1 X PBS and incubated in DAPI (1µg/ml) for 10 min at RT and then washed with 1X PBS and stored at 4°C until confocal/fluorescence imaging. (53) The mitotic phenotypes were quantified using 3D reconstruction of the confocal images using the IMARIS software (54). Rabbit antibodies: anti-γ tubulin (1:2000) (Thermo#PA5-34815), Mouse antibodies: anti-α tubulin (1:100) (CST#3873).

### Akt Activator assay and acridine orange staining

DIS3L2 morpholino injected embryos at the shield stage (6hpf) were exposed to 1uM of Akt activator SC79 (Tocris, # 4635). Exposure to 0.5% DMSO (vehicle) was used as a control. The control and DIS3L2 injected embryos treated with SC79 were phenotyped at 24hpf and stained with acridine orange to detect the apoptotic cells in the embryos. For acridine orange staining, the embryos were dechorinated, and stained with 5mg/L AO for 1 hour in the dark followed by washing with embryo water. Then the embryos were anesthetized with tricaine and were photographed by a Zeiss AxioZoom.V16 microscope, the bright green dots in the embryos indicate apoptotic cells. The number of apoptotic cells in the head region was counted using Image J software (27,48,51).

### Skeletal preparations by Alcian blue and Alizarin red staining

Wild-type control and morphants 5-day-old larvae were fixed in 4% PFA/PBS overnight at 4^0^C. The embryos were washed with 1XPBS and stored overnight in 100% methanol at - 20^0^C. The following staining solutions were prepared - Solution A (0.04% Alcian blue, 125mM MgCl2 in 70% ethanol) and Solution B (1.5% Alizarin Red S). 1ml of solution A and 10ul of solution B were added to the larvae and kept overnight at room temperature with constant rocking. The staining solution was removed and the larvae were washed two times with water, followed by bleach solution for 1 hour at room temperature and a gradation of Glycerol-KOH. The embryos were then stored in 50% glycerol/ 0.25% KOH for 2 hours at room temperature. The stained larvae were then imaged using a Zeiss Stemi508 stereomicroscope (51).

## Results

### *Dis3l2* depletion results in neurodevelopmental defects

To ascertain the developmental functions of DIS3L2, we first examined the gene expression profile of *dis3l2* across developmental stages of zebrafish (Fig. 1). We observed *dis3l2* transcript expression in the cleavage stages (2 cell), gastrula stage (50% epiboly), somitogenesis (6-18 somites) and in 24hpf by RT-PCR analysis (Fig. 1A). We also examined the spatiotemporal expression pattern of *dis3l2* in early development, somite stages and in 24hpf embryos by *in situ* hybridization (Fig. 1A). We observed that *dis3l2* is expressed ubiquitously during early embryogenesis and the expression is restricted to the anterior part of the embryo, particularly the brain (red asterisk, Fig. 1A) and in the eye in 24hpf embryos (red arrowhead, Fig. 1A). At the cellular level, DIS3L2 localizes to the nucleus (white asterisk, Fig.1B) and cortex in the blastula stage (256 cell stage) (white arrowhead, Fig. 1B). To probe the function of DIS3L2 in embryogenesis, we used the antisense morpholino-based gene knockdown approach (Fig.1C). The *dis3l2*-depleted embryos showed a range of phenotypes. About 18% of morpholino-injected embryos showed normal morphology and were termed as P0 (Fig. 1C). About 58% of the *dis3l2*-depleted embryos that exhibited neurodevelopmental defects, poorly formed brains, and disrupted brain architecture were classified as P1 (red asterisk, Fig. 1C). The severe phenotype termed P2 (22%), showed acute neurodevelopmental abnormalities (double red asterisk, Fig.1C) and a complete lack of midbrain-hindbrain boundary (MHB) (red arrowhead, Fig. 1C). The severity of the phenotypes P0, P1, and P2 was observed in a morpholino concentration-dependent manner, as the number of P1 and P2 phenotypes increased in the embryos injected with higher morpholino concentration (Fig. 1D). Western blot analysis of control and *dis3l2* morpholino-injected embryonic lysates showed a significant reduction in DIS3L2 protein levels, thus confirming the DIS3L2 specific depletion (Fig. 1E). Further, the *dis3l2*-depleted embryos showed an overall increase in mortality by 20% (Fig. 1F). The morphants that survived to larval stages, exhibited craniofacial defects, as shown by Alcian blue and Alizarin red S staining of bone and cartilage (red asterisk, Fig. 1G, H). The morphants larvae also exhibited cardiac edema (red arrowhead, Fig. 1G). These observations suggest that *dis3l2* mRNA is maternally inherited in the embryo and plays a key role in early brain development.

**Figure 1:**
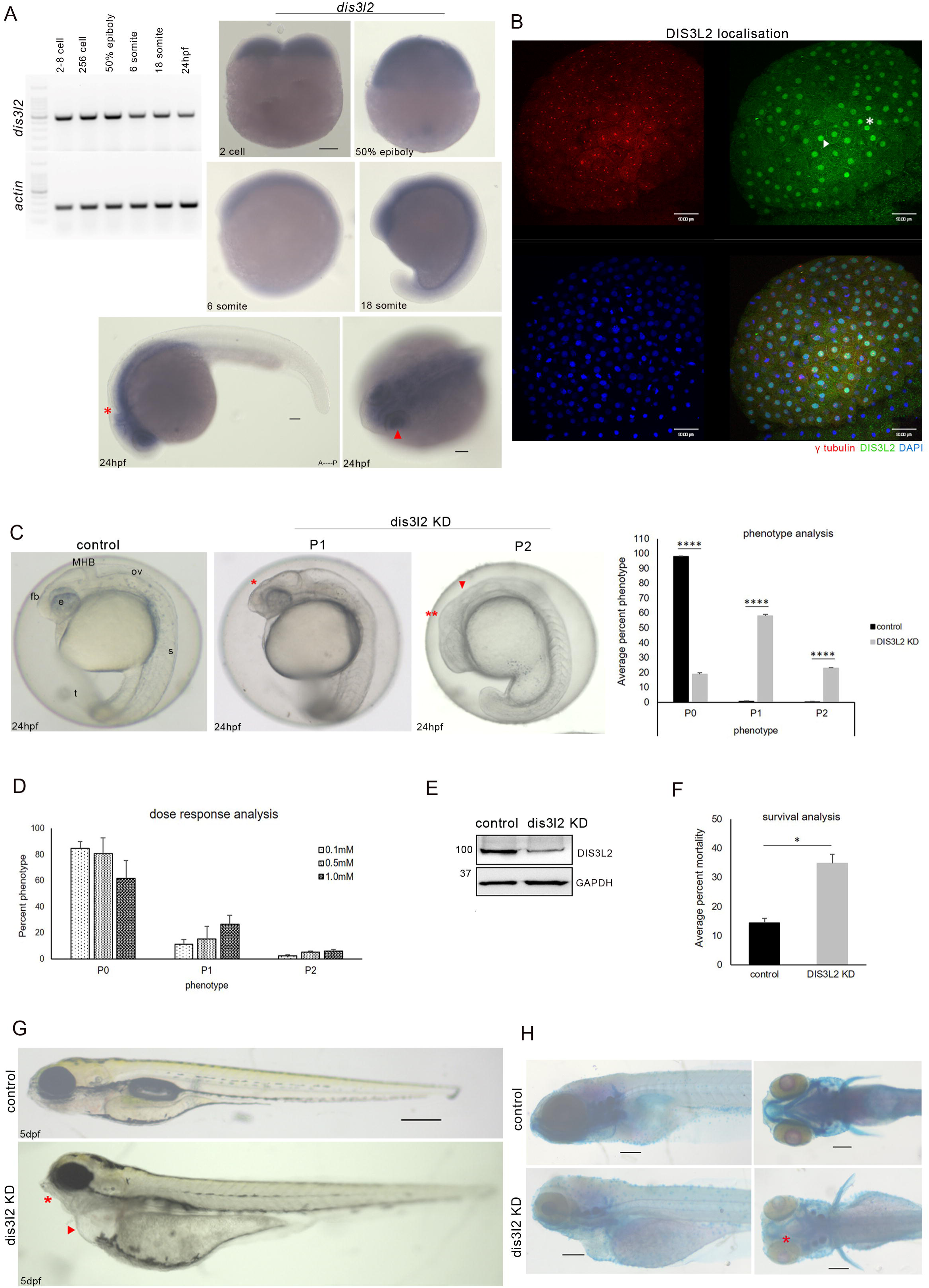
*Dis3l2* depletion results in neurodevelopmental defects. A: RT-PCR analysis of *dis3l2* transcripts in different developmental stages. Zebrafish *actin* was used as the loading control (*left*). Whole-mount RNA *in situ* hybridization showing *dis3l2* transcripts during early development. Scale bars, 100µm (*right*). B: Sum projection confocal image of 256 cell stage control embryos showing endogenous DIS3L2 localization. Blue/DAPI - DNA, green – DIS3L2, red – γ tubulin. Scale bars, 50µm. C: Gross morphological analysis of *dis3l2* depleted embryos by morpholino in comparison to control embryos at 24hpf showing forebrain defects (red asterisks), midbrain-hindbrain boundary defects (red arrowhead) Quantification of percent phenotype of 1mM *dis3l2* morphants as compared to stage-matched control. Data are shown as mean+ SEM, *p<0.05, **p<0.01, ***p<0.001. n=3 for each experiment. D: Percent phenotype of *dis3l2* morphants in different concentrations of morpholino. E: Western blot analysis showing DIS3L2 levels by western blot in *dis3l2* morpholino injected morphants. GAPDH was used as the loading control. F: Percent mortality of *dis3l2* morphants with respect to control at 24hpf. G: Bright field image of *dis3l2* depleted larvae as compared to control larvae at 5dpf. H: Skeletal preparations of *dis3l2* morphants by bone and cartilage staining using Alizarin red S and Alcian blue respectively. Scale bars 50µm.

### *Dis3l2* is crucial for the specification of neural crest cells

To investigate the developmental basis of neural crest-derived craniofacial abnormalities in DIS3L2-depleted embryos (Fig.1G, H), we examined the expression dynamics of neural crest specifier genes *foxd3*, *crestin*, *twist1a* and *sox10* (Fig.2) (7). *Foxd3* is essential for survival of hindbrain neural crest cells and plays a key role in specification of the premigratory neural crest cells into the neuronal, glial or cartilage fate by inducing expression of lineage specific transcription factors (8,9). *Foxd3* is downregulated in the cranial neural crest of the morphants (red asterisk, Fig.4A). Intriguingly, *foxd3* expression is retained in the caudal neural crest and somites. On the other hand, *crestin* expression in the post otic trunk neural crest increased in the DIS3L2 morphants (red arrow, Fig. 4B). *Crestin* is a pan neural crest marker which is expressed after the neural crest cells are specified until they undergo overt differentiation (10–12). *Twist1a* is a neural crest specifier expressed in the premigratory cranial neural crest in the diencephalon and is essential for the migration and differentiation (13). *Sox10* is also expressed in undifferentiated neural crest cells and is required for neural crest specification, migration, and survival (12–16). The DIS3L2 morphants showed downregulation of both *twist1a* and *sox10* (red asterisks, Fig.4C, D). The loss of *sox10* expression results in cell death and failure to undergo differentiation to give rise to the neural crest-derived craniofacial skeleton (17,18). The down regulation of these specifiers attribute the craniofacial anomalies observed in the dis3l2 morphants. We also examined the neural tube patterning and differentiation genes like *pax2a* and *fgf8*. *Pax2a* is expressed in the optic stalk, MHB, eye, otic placode and promotes otic differentiation (19–22). *Fgf8* is also expressed in the MHB and eye (23,24). *Pax2a* (red arrow, Fig.4E) and *fgf8* (red asterisk, Fig.4F) are downregulated in *dis3l2* morphants. The loss of structures in the midbrain and anterior hindbrain in the *dis3l2* morphants may be attributed to the reduced expression of *fgf8* and loss of inductive signals from the *Fgf8* expressing hindbrain tissue. Thus, DIS3L2 is essential to initiate the neural differentiation program, and its precocious depletion results in the loss of differentiation-inducing signals in the embryo.

**Figure 2:**
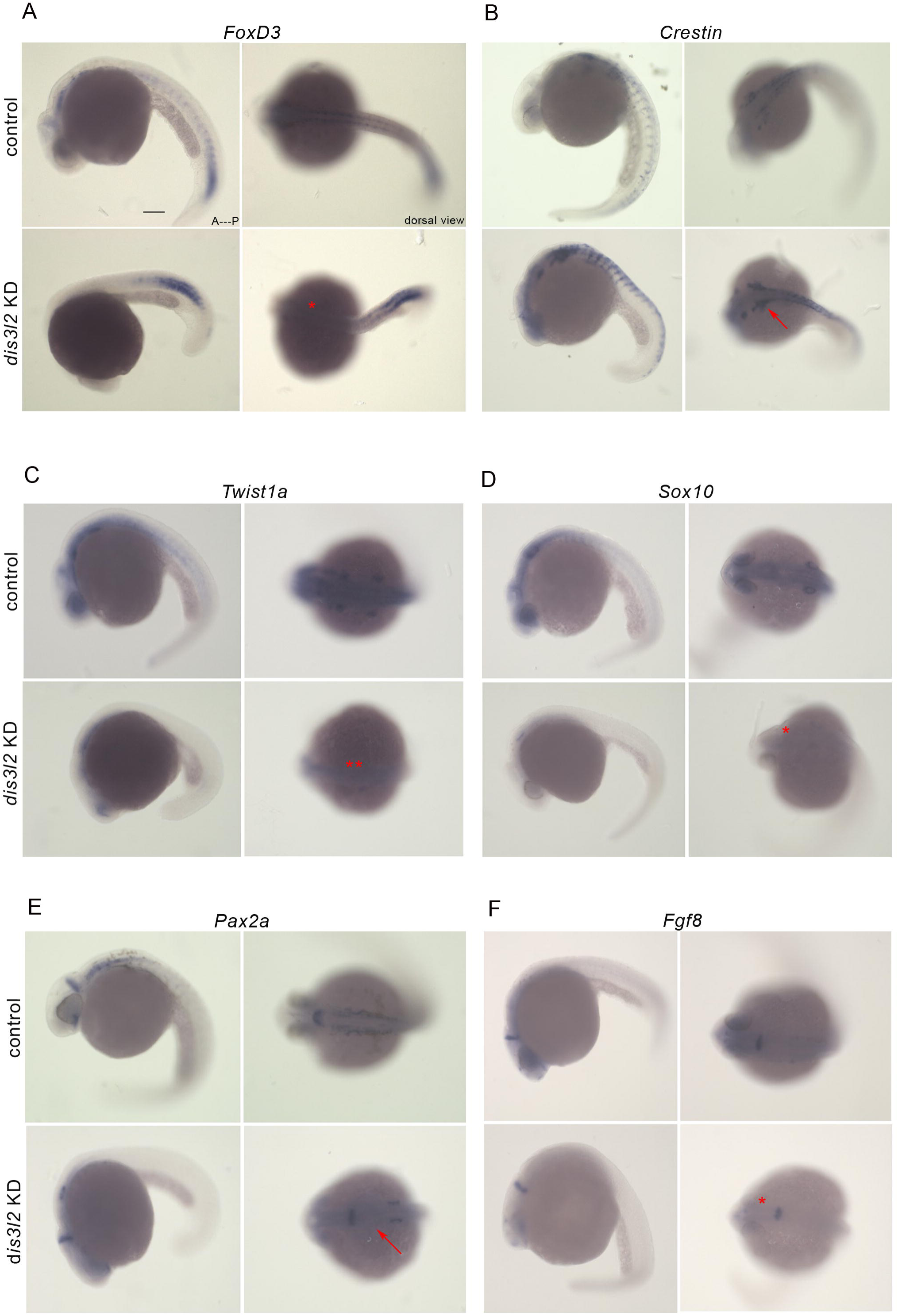
*Dis3l2* is crucial for the specification of neural crest cells. A, B: Gene expression analysis of *foxd3* (red asterisk) and *crestin* (red arrow) in *dis3l2* morphants with respect to control. C, D: Expression of *twist1a* (double red asterisks) and *sox10* (single red asterisk) in *dis3l2* morphants as compared to control. E, F: Gene expression of *pax2a* (red arrow) and *fgf8* in *dis3l2* morphants and control. Scale bars, 100µm.

### DIS3L2 is essential for neural progenitor survival

In addition to craniofacial defects, the DIS3L2 morphant P1 larvae also exhibited smaller head size at 5dpf (Fig.3A). To determine the cellular basis of the smaller head phenotype, we examined the status of cell death in the DIS3L2 P1 morphants by acridine orange staining (Fig. 3B). Intriguingly, we observed a massive surge of apoptosis in the developing brain and neural tube of the *dis3l2* morphants, indicating that DIS3L2 associated head size reduction might be due to increased apoptosis (white asterisk, Fig. 3B) and reduced expression of neural tube markers such as *pax2a* (Fig.4E, F). As the intrinsic apoptosis pathway responds to developmental cues, we first examined the pro-apoptotic Bcl2 family, initiator protease CASPASE 9, and associated effector CASPASE 3 (25–27). Remarkably, the expression of pro-apoptotic Bcl2 members *BIM*, *puma,* and *foxo* increased in the DIS3L2 morphants (Fig. 3C). However, the pro-survival *Bcl2* levels remain unchanged in the DIS3L2 morphants (Fig3C). *BIM*, *puma*, and *foxo* activate CASPASE9 and CASPASE3 upon apoptotic stimulation (27–30). We also observed an increase in CASPASE3 and CASPASE9 levels in the DIS3L2-depleted embryonic lysates (Fig. 3D). Elevated PUMA levels are known to promote neuronal cell death by activating both intrinsic and extrinsic apoptotic pathways in the zebrafish embryonic neural tissue (27,31,32). Thus, the over-expression of these proapoptotic genes may be the underlying basis of increased cell death in the *dis3l2* morphants. And the depletion of DIS3L2, induces pro-death signals, leading to activation of these BH3-only class of Bcl2 proteins and elevated CASPASE levels. Reports show that transcriptional repression of BIM is a key cell survival signal and its activity is regulated through phosphorylation by Akt signaling (27,29,30). To further probe into the mechanism of DIS3L2-mediated apoptosis, we therefore examined the Akt signaling pathway. Akt signaling has been shown to regulate brain size in the zebrafish embryo and reduced Akt signaling results in a reduction in brain size. Further, over-expression of the constitutive form of Akt inhibits neuronal apoptosis (7,19,33–35). We used immunoblotting assay to determine the phosphorylation levels of Akt in control and DIS3L2-depleted embryos. AKT (pAKT-Thr308) phosphorylation levels decreased in DIS3L2 morphants, while the total AKT remained unchanged. (Fig.3E). Phosphorylation at Thr308 is critical for the activation loop of the kinase by protein kinase PDK1 and activation of AKT signaling. Akt in turn, phosphorylates GSK3β, which also affects cell survival, learning, and memory in the brain (33,36). As the phosphorylation status of GSK3β at ser9 position dictates its kinase activity and the phosphorylated form is pro-survival, we examined the phosphorylated GSK3β levels in DIS3L2-depleted embryonic lysates. We observed that phosphorylated GSK3β (pGSK3β Ser9) was reduced in DIS3L2 morphants and the total GSK3β levels did not change. (Fig. 3E). Thus, decreased GSK3β levels are concomitant with a significant decrease in Akt signaling in the DIS3L2 morphants. To ascertain direct DIS3L2-mediated regulation of apoptosis by Akt signaling, we treated the DIS3L2 morphants to Akt activator SC79. SC79 binds to Akt, enhances its phosphorylation, and activates cytosolic Akt (37,38). The embryos treated with the Akt activator showed a significant reduction in DIS3L2 phenotypes P1 and P2 (Fig. 3F) and apoptosis, marked by acridine orange staining (white arrowhead, Fig. 3G). These data strongly suggest that *DIS3L2* is essential for neural progenitor survival and regulating apoptosis during CNS morphogenesis via Akt signaling.

**Figure 3:**
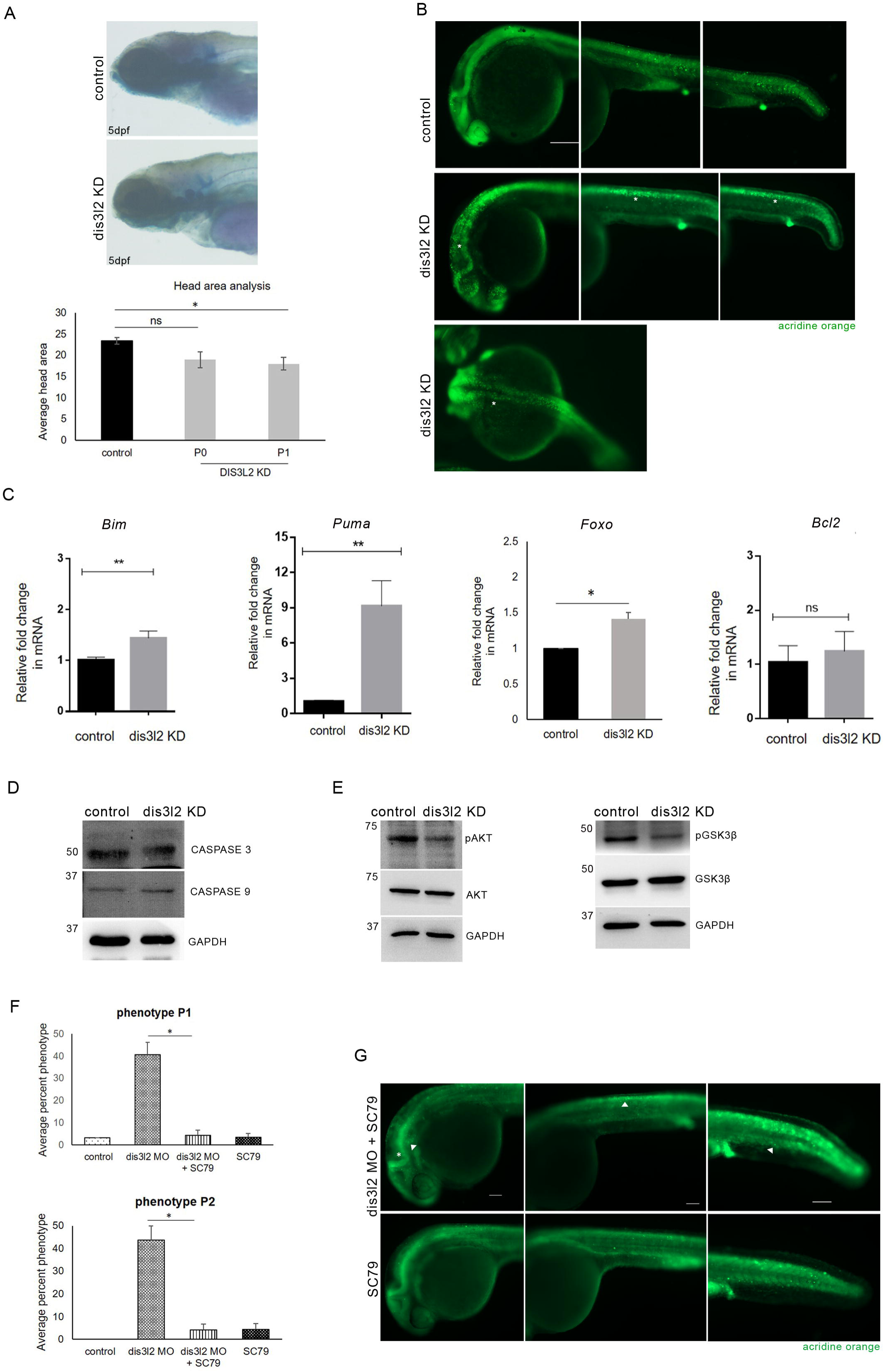
*Dis3l2* is essential for neural progenitor survival. A: Skeletal preparation and head measurements of *dis3l2* depleted larvae showing smaller head area with respect to control. B: Cell death analysis in the *dis3l2* morphants by acridine orange staining. Acridine orange positive cells (white asterisk) shown in the head. Scale bars, 200µm. C: Relative expression of Bcl2 family members – *bim*, *puma, foxo,* and *bcl2* in *dis3l2* morphants. Data are shown as mean+ SEM, *p<0.05, **p<0.01, ***p<0.001. n=3 for each experiment. D: Western blot analysis to show CASPASE3 levels in morphants embryos. GAPDH was used as loading control. E: Western blot analysis showing phosphorylated AKT with respect to total AKT levels and phosphorylated GSK3β with respect to total GSK3β levels in *dis3l2* depleted embryos. GAPDH was used as loading control. F: Rescue experiments showing phenotype P1 and P2 upon treatment with Akt activator SC79 in *dis3l2* morpholino injected embryos. G: Cell death analysis by acridine orange staining in *dis3l2* morpholino injected embryos treated with SC79 and WT embryos treated with SC79.

### DIS3L2 is required for early embryonic mitoses

As DIS3L2 plays an important role in regulating apoptosis in the developing CNS, we wanted to investigate its possible functions in regulating mitosis and the cell cycle (Fig. 4). Indeed, *dis3l2* morphants showed a myriad of mitotic defects during the early blastula stage (Fig. 4A). The morphants exhibited elongated spindles (white dotted line, Fig. 4A) and chromosome mis-congression at metaphase (Fig. 3A, B, C), suggesting that DIS3L2 is essential to maintain spindle length and aids in chromosome congression during metaphase. The morphants also exhibited unfocused spindle poles with multiple γ-tubulin positive foci during interphase and mitosis (white asterisk, Fig. 3A, B, C). The morphant embryos also showed an increased cytokinetic index, indicating its role in cytokinetic progression (white arrowhead, Fig.3B, C). Hence, our results show that DIS3L2 is essential for mitotic spindle organization, spindle pole integrity and mitotic conformity during early embryonic divisions. We also examined the effect of DIS3L2 depletion on cell cycle regulators p21 and p27. These are key cell cycle inhibitors that suppress cell cycle progression by inhibiting cyclin-dependent kinases (CDKs) and arresting cells in the G1 phase. p21 is also a cellular marker of senescence, which is characterized by cell cycle arrest and loss of function (39–42). The morphants showed an overall upregulation of *p21*, *p27a,* and *cyclinD1* (Fig. 4D). The upregulation of *p21* and *cyclinD* transcripts may be the underlying cause of impaired cell cycle progression, increased cell death, and growth retardation in the *dis3l2* morphants. Further, cell cycle progression is tightly regulated by cyclins and CDKs complexes (41,43,44). The mitotic CDK, CDK1, a downstream target of p21, is essential for cell division and recently reported for cell survival. CDK1 promotes cell death by activating Bcl2 family members like BAD and phosphorylates MCL1, suppressing its anti-apoptotic functions (43–45). Hence, the upregulation of CDK1 and pro-apoptotic *BIM* and *Puma* transcripts strongly corroborates with the surge of apoptosis in the *dis3l2* morphants (Fig. 4E). Also, the Mek/Erk (MAPK) pathway is a key signaling cascade that regulates cell division, differentiation, and apoptosis. MEK signaling is also essential for cranial neural crest specification, formation of Meckel’s cartilage, ceratohyal cartilage, and neural crest-derived craniofacial skeletogenesis (46–49). Interestingly, we observed a downregulation of MEK1 (MAP2K1) in the DIS3L2 morphants (Fig.4E), which also contributes to the observed craniofacial defects (Fig.1G, H). Together, we demonstrate that DIS3L2 plays a key role in cell cycle regulation and mitotic progression during early embryogenesis.

**Figure 4:**
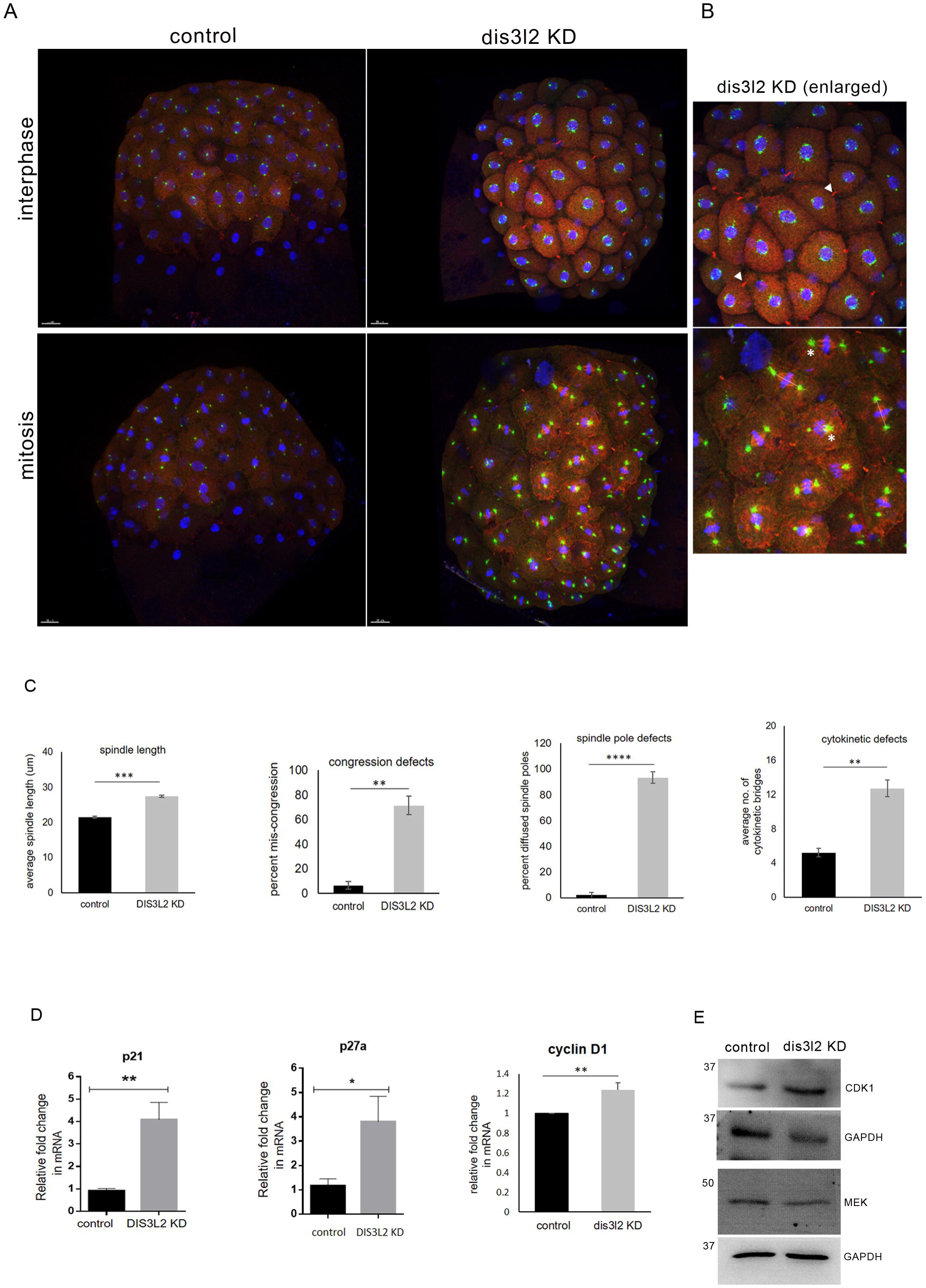
DIS3L2 is required for early embryonic mitoses. A: Sum projection confocal image of 256 cell stage control and *dis3l2* morphants embryos showing mitotic defects. Blue/DAPI - DNA, red – alpha-tubulin, green – γ tubulin. B: Enlarged sum projection confocal image of *dis3l2* morphants showing cytokinetic index (white arrowhead), spindle length (white dotted line), and spindle poles (white asterisks) during early mitoses. Scale bars, 50µm. C: Quantification of spindle length (white dotted line), chromosome congression, spindle pole (white asterisk), and cytokinesis (white arrowhead) in the *dis3l2* morphants as compared to control. Scale bars, 50µm. All data are shown as mean + SEM. *p<0.05, **p<0.01, ***p<0.001. n=3 for each experiment. D: Relative expression of cell cycle regulators – *p21*, *p27,* and *cyclinD1* in *dis3l2* morphants. Data are shown as mean+ SEM, *p<0.05, **p<0.01, ***p<0.001. n=3 for each experiment. E: Western blot analysis showing CDK1 and MEK1 levels in *dis3l2* depleted embryos as compared to control. GAPDH was used as the loading control.

## Discussion

DIS3L2 is an exoribonuclease that can degrade single and double-stranded RNA substrates (1,3,50). *In vitro* studies show that DIS3L2 negatively regulates cell cycle progression and its depletion results in mitotic aberrations and tumor growth (4). A recent report shows that DIS3L2 regulates cell proliferation and tissue growth by Idgf2 signaling pathway coupled with increased Akt phosphorylation levels (6). Yet, the molecular mechanism of DIS3L2 function in cell proliferation and differentiation during vertebrate embryogenesis remains elusive. This study delineates novel developmental functions of DIS3L2 in early embryonic mitoses and neurodevelopment. Depletion of DIS3L2 results in upregulation of cell cycle regulators p21, p27 and cyclinD1 and mitotic aberrations such as elongated metaphase spindles, diffused spindle poles, chromosome mis-congression, and cytokinetic defects. These results show that DIS3L2 is required for mitotic progression during early embryonic development and is essential for proper chromosome congression, maintenance of spindle pole integrity, and cytokinesis.

In the context of embryogenesis, we show that DIS3L2 is maternally expressed during early embryonic stages and plays a key role in brain morphogenesis and architecture. Our study reveals that *dis3l2* plays a key role in cranial neural crest specification. The reduced expression of neural crest specifier genes *foxd3*, *twist1a,* and *sox10* in morphants indicates that DIS3L2 plays a key role in specifying progenitor cell fate, migration, and differentiation of neural crest cells. The reduced expression of neural crest specifiers like *sox10* and decreased MEK signaling contribute to observed craniofacial skeletal abnormalities. Thus, *dis3l2* is critical for the fate determination of neural crest and its derivatives. Further, *dis3l2* depletion resulted in massive cell death and upregulation of pro-apoptotic genes *bim*, *puma,* and *foxo3a* in the embryonic neural tissue. Previous studies show that neuronal survival and growth in the developing embryo are attributed to the phosphoinositide 3 kinase (PI3K) / Akt kinase cascade. PI3K stimulates the activation of Akt which in turn, regulates cell size, proliferation, and survival. Our results demonstrate that DIS3L2 is essential for neural progenitor survival by regulating the Akt signaling. DIS3L2 depletion showed down-regulation of Akt-GSK3β signaling, and the administration of Akt activator SC79 rescued cell death. Akt signaling also regulates neural crest gene expression in vivo, which further potentiates our DIS3L2-mediated Akt and neural crest specifier genes down-regulation (7). These findings are summarised in Fig.5.

**Figure 5:**
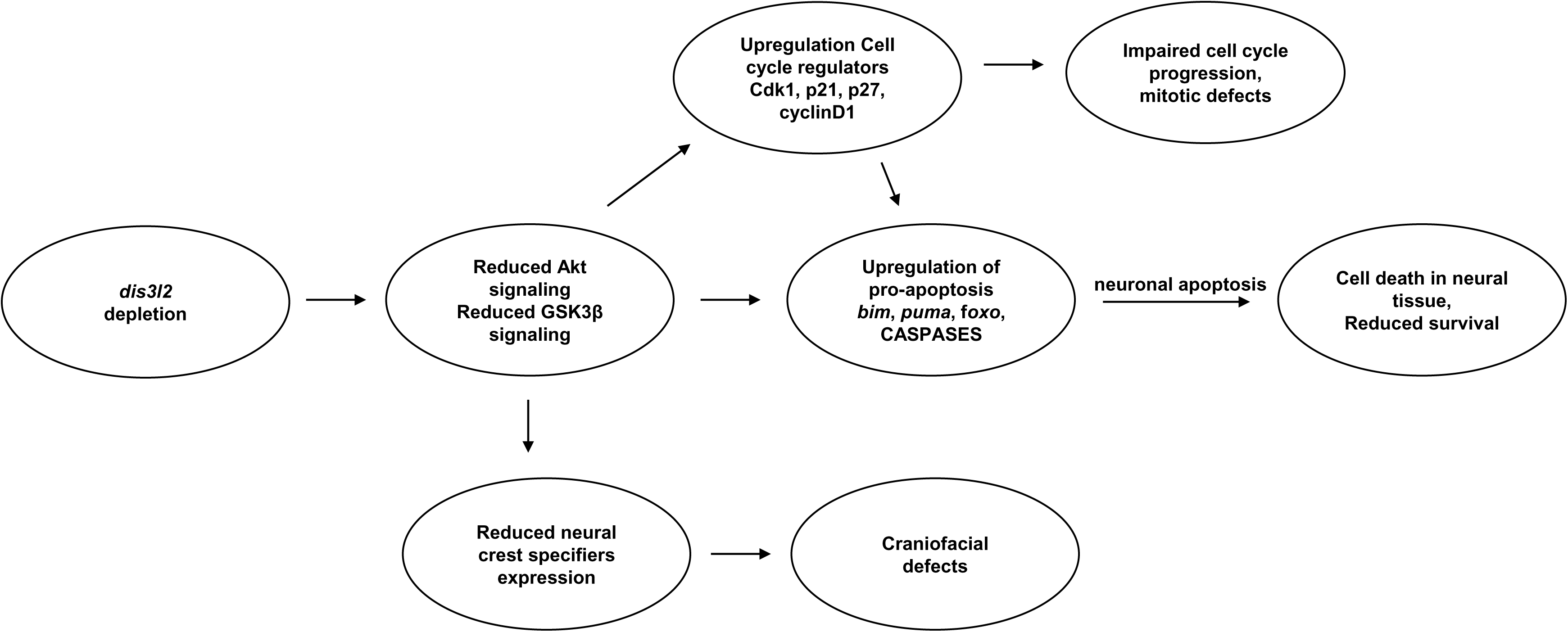
Summary of developmental functions of *dis3l2*. Schematic representation to synopsize role of *dis3l2* in early embryogenesis. *Dis3l2* depletion results in reduced Akt and GSK3β signaling, resulting in upregulation of apoptosis pathways involving *bim*, *puma*, *foxo* and effector CASPASES. This results in massive surge of apoptosis in the embryonic neural tissue. Concomitantly, *dis3l2* depletion also results in upregulation of cell cycle regulators such as CDK1, p21 and p27, which in turn further upregulate the apoptotic mechanisms. Hence, the embryos show myriad of mitotic defects during early embryogenesis. Additionally, the *dis3l2* morphants exhibit reduced expression of neural crest specifier genes, resulting in severe craniofacial anomalies at larval stages.

### Limitations of the study

The current study is limited to cell fate specification and survival of neural progenitor cells in the embryonic CNS. The DIS3L2-mediated mechanism of cell survival in non-CNS tissues such as mesodermal tissue has not been addressed. It is imperative to also determine how DIS3L2 recruits its targets to perform its diverse cellular functions. Lastly, the mechanism by which DIS3L2 interacts with the mitotic machinery remains to be elucidated.

## Conclusions

Our study provides fresh perspectives on DIS3L2-mediated neuronal apoptosis and its neurodevelopmental functions, particularly in cranial neural crest specification, patterning, and survival. DIS3L2 mutations are associated with Perlman syndrome and Wilm’s tumor (1,3,50). Future studies to establish a DIS3L2-based Perlman syndrome model using zebrafish embryos will provide mechanistic insights into the pathophysiology and cellular basis of the affected neural tissue. Delineating such molecular mechanisms governing neuronal apoptosis and attainment of normal brain size will also provide a better understanding of neurological diseases like Alzheimer’s and Parkinson’s disease.

## Supporting information

supplementary_westernblots

## List of abbreviations

DIS3L2: DIS3 like 3’-5’ exoribonuclease 2
Hpf: hours post fertilization
Dpf: days post fertilization
Akt: Protein Kinase B/ PKB
GSK3β: Glycogen synthase kinase 3β

## Declarations

### Ethics approval and consent to participate

Not applicable.

### Consent for publication

All authors have read the manuscript and given consent for publication.

### Availability of data and materials

Not applicable.

### Author contributions

SD and TP contributed equally to this work. SD performed all the zebrafish experiments, analyzed the data, and assisted in manuscript writing and figure preparation. TP performed all the western blot assays, Akt activator assays and assisted in figure preparation. MK developed the concept and experiments, supervised and mentored the entire study, prepared the figures, and wrote the manuscript.

## Acknowledgements

The authors thank the members of the MK lab for critical reading of the manuscript.

## Legends

**Fig. S1**: List of primers used in this study.

**Fig. S2:** Raw, uncropped images of the western blots shown in this study.

## References

1. Hunter RW, Liu Y, Manjunath H, Acharya A, Jones BT, Zhang H, et al. Loss of Dis3l2 partially phenocopies perlman syndrome in mice and results in upregulation of Igf2 in nephron progenitor cells. Genes Dev. 2018;32(13–14):903–8.

2. Thomas MP, Liu X, Whangbo J, McCrossan G, Sanborn KB, Basar E, et al. Apoptosis Triggers Specific, Rapid, and Global mRNA Decay with 3’ Uridylated Intermediates Degraded by DIS3L2. Cell Rep [Internet]. 2015;11(7):1079–89. Available from: 10.1016/j.celrep.2015.04.026

3. Luan S, Luo J, Liu H, Li Z. Regulation of RNA decay and cellular function by 3′-5′ exoribonuclease DIS3L2. RNA Biol [Internet]. 2019;16(2):160–5. Available from: 10.1080/15476286.2018.1564466

4. Astuti D, Morris MR, Cooper WN, Staals RHJ, Wake NC, Fews GA, et al. Germline mutations in DIS3L2 cause the Perlman syndrome of overgrowth and Wilms tumor susceptibility. Nat Genet. 2012;44(3):277–84.

5. Towler BP, Jones CI, Harper KL, Waldron JA, Newbury SF. A novel role for the 3′-5′ exoribonuclease Dis3L2 in controlling cell proliferation and tissue growth. RNA Biol. 2016;13(12):1286–99.

6. Towler BP, Pashler AL, Haime HJ, Przybyl KM, Viegas SC, Matos RG, et al. Dis3L2 regulates cell proliferation and tissue growth though a conserved mechanism. PLoS Genet [Internet]. 2020;16(12 December):1–29. Available from: 10.1371/journal.pgen.1009297

7. Ciarlo C, Kaufman CK, Kinikoglu B, Michael J, Yang S, D’Amato C, et al. A chemical screen in zebrafish embryonic cells establishes that Akt activation is required for neural crest development. Elife. 2017;6:1–26.

8. Stewart RA, Arduini BL, Berghmans S, George RE, Kanki JP, Henion PD, et al. Zebrafish foxd3 is selectively required for neural crest specification, migration and survival. Dev Biol. 2006;292(1):174–88.

9. Lister JA, Cooper C, Nguyen K, Modrell M, Grant K, Raible DW. Zebrafish Foxd3 is required for development of a subset of neural crest derivatives. Dev Biol [Internet]. 2006;290(1):92–104. Available from: 10.1016/j.ydbio.2005.11.014

10. Luo R, An M, Arduini BL, Henion PD. Specific pan-neural crest expression of zebrafish crestin throughout embryonic development. Dev Dyn. 2001;220(2):169–74.

11. Corallo D, Candiani S, Ori M, Aveic S, Tonini GP. The zebrafish as a model for studying neuroblastoma. Cancer Cell Int. 2016;16(1):1–9.

12. Asad Z, Pandey A, Babu A, Sun Y, Shevade K, Kapoor S, et al. Rescue of neural crest-derived phenotypes in a zebrafish CHARGE model by Sox10 downregulation. Hum Mol Genet. 2016 Aug 15;25(16):3539–54.

13. Yeo GH, Cheah FSH, Jabs EW, Chong SS. Zebrafish twist1 is expressed in craniofacial, vertebral, and renal precursors. Dev Genes Evol. 2007 Dec;217(11– 12):783–9.

14. Dutton KA. Zebrafish sox10 mutant [Internet]. 2001. Available from: www.hgmp.mrc.ac.uk/Registered/Option/mrtrans.html

15. Drerup CM, Wiora HM, Topczewski J, Morris JA. Disc1 regulates foxd3 and sox10 expression, affecting neural crest migration and differentiation. Development. 2009 Aug 1;136(15):2623–32.

16. Nagao Y, Takada H, Miyadai M, Adachi T, Seki R, Kamei Y, et al. Distinct interactions of Sox5 and Sox10 in fate specification of pigment cells in medaka and zebrafish. PLoS Genet. 2018 Apr 1;14(4 April).

17. Carney TJ, Dutton KA, Greenhill E, Delfino-Machín M, Dufourcq P, Blader P, et al. A direct role for Sox10 in specification of neural crest-derived sensory neurons. Development. 2006 Dec;133(23):4619–30.

18. Delfino-Machín M, Madelaine R, Busolin G, Nikaido M, Colanesi S, Camargo-Sosa K, et al. Sox10 contributes to the balance of fate choice in dorsal root ganglion progenitors. PLoS One. 2017 Mar 1;12(3).

19. Chen S, Liu Y, Rong X, Li Y, Zhou J, Lu L. Neuroprotective role of the PI3 Kinase/Akt signaling pathway in Zebrafish. Front Endocrinol (Lausanne). 2017;8(FEB).

20. Hans S, Liu D, Westerfield M. Pax8 and Pax2a function synergistically in otic specification, downstream of the Foxi1 and Dlx3b transcription factors. Development. 2004 Oct;131(20):5091–102.

21. Lusk S, Kwan KM. Pax2a, but not pax2b, influences cell survival and periocular mesenchyme localization to facilitate zebrafish optic fissure closure. Dev Dyn. 2022 Apr 1;251(4):625–44.

22. Tan AL, Christensen SE, Baker AK, Riley BB. Fgf, Hh, and pax2a differentially regulate expression of pax5 and pou3f3b in vestibular and auditory maculae in the zebrafish otic vesicle. Dev Dyn. 2023 Oct 1;252(10):1269–79.

23. Carraco G, Martins-Jesus AP, Andrade RP. The vertebrate Embryo Clock: Common players dancing to a different beat. Vol. 10, Frontiers in Cell and Developmental Biology. Frontiers Media S.A.; 2022.

24. Wiellette EL, Sive H. Early Requirement for fgf8 Function during Hindbrain Pattern Formation in Zebrafish. Dev Dyn. 2004 Feb;229(2):393–9.

25. Eimon PM, Kratz E, Varfolomeev E, Hymowitz SG, Stern H, Zha J, et al. Delineation of the cell-extrinsic apoptosis pathway in the zebrafish. Cell Death Differ. 2006;13(10):1619–30.

26. Yabu T, Kishi S, Okazaki T, Yamashita M. Characterization of zebrafish caspase-3 and induction of apoptosis through ceramide generation in fish fathead minnow tailbud cells and zebrafish embryo. Biochem J. 2001;360(1):39–47.

27. Eimon PM, Ashkenazi A. The zebrafish as a model organism for the study of apoptosis. Apoptosis. 2010;15(3):331–49.

28. Kratz E, Eimon PM, Mukhyala K, Stern H, Zha J, Strasser A, et al. Functional characterization of the Bcl-2 gene family in the zebrafish. Cell Death Differ. 2006;13(10):1631–40.

29. Jette CA, Flanagan AM, Ryan J, Pyati UJ, Carbonneau S, Stewart RA, et al. BIM and other BCL-2 family proteins exhibit cross-species conservation of function between zebrafish and mammals. Cell Death Differ. 2008;15(6):1063–72.

30. Bodaar K, Colon D, Pyati U, Stevenson KE, Qi J. in high-risk T-cell acute lymphoblastic leukemia. 2015;28(9):1819–27.

31. Rodríguez-Aznar E, Nieto MA. Repression of Puma by Scratch2 is required for neuronal survival during embryonic development. Cell Death Differ. 2011;18(7):1196–207.

32. Toruno C, Carbonneau S, Stewart RA, Jette C. Interdependence of bad and puma during ionizing-radiation-induced apoptosis. PLoS One. 2014;9(2).

33. Moore NS, Mans RA, McCauley MK, Allgood CS, Barksdale KA. Critical effects on akt signaling in adult zebrafish brain following alterations in light exposure. Cells. 2021;10(3):1–11.

34. Thiagarajan SK, Mok SY, Ogawa S, Parhar IS, Tang PY. Receptor-Mediated AKT/PI3K Signalling and Behavioural Alterations in Zebrafish Larvae Reveal Association between Schizophrenia and Opioid Use Disorder. Int J Mol Sci. 2022;23(9).

35. Cheng YC, Hsieh FY, Chiang MC, Scotting PJ, Shih HY, Lin SJ, et al. Akt1 Mediates Neuronal Differentiation in Zebrafish via a Reciprocal Interaction with Notch Signaling. PLoS One. 2013;8(1):1–12.

36. Finkielsztein A, Kelly GM. Altering PI3K—Akt signalling in zebrafish embryos affects PTEN phosphorylation and gastrulation. Biol Cell. 2009;101(11):661–82.

37. Jo H, Mondal S, Tan D, Nagata E, Takizawa S, Sharma AK, et al. Small molecule-induced cytosolic activation of protein kinase Akt rescues ischemia-elicited neuronal death. Proc Natl Acad Sci U S A. 2012;109(26):10581–6.

38. Luan Q, Pan L, He D, Gong X, Zhou H. SC79, the AKT activator protects cerebral ischemia in a Rat Model of Ischemia/Reperfusion Injury. Med Sci Monit. 2018;24:5391–7.

39. Masai I, Yamaguchi M, Tonou-Fujimori N, Komori A, Okamoto H. The hedgehog- PKA pathway regulates two distinct steps of the differentiation of retinal ganglion cells: The cell-cycle exit of retinoblasts and their neuronal maturation. Development. 2005;132(7):1539–53.

40. Langheinrich U, Hennen E, Stott G, Vacun G. Zebrafish as a model organism for the identification and characterization of drugs and genes affecting p53 signaling. Curr Biol. 2002;12(23):2023–8.

41. Lataster L, Huber HM, Böttcher C, Föller S, Radziwill G. Cell Cycle Control by Optogenetically Regulated Cell Cycle. Biology (Basel). 2023;12(1194):1194.

42. Laranjeiro R, Tamai TK, Letton W, Hamilton N, Whitmore D. Circadian Clock Synchronization of the Cell Cycle in Zebrafish Occurs through a Gating Mechanism Rather Than a Period-phase Locking Process. J Biol Rhythms. 2018;33(2):137–50.

43. Jin M, Li J, Hu R, Xu B, Huang G, Huang W, et al. Cyclin A2/cyclin-dependent kinase 1-dependent phosphorylation of Top2a is required for S phase entry during retinal development in zebrafish. J Genet Genomics [Internet]. 2021;48(1):63–74. Available from: 10.1016/j.jgg.2021.01.001

44. Massacci G, Perfetto L, Sacco F. The Cyclin-dependent kinase 1: more than a cell cycle regulator. Br J Cancer. 2023;129(11):1707–16.

45. Boos A, Gahr BM, Park DD, Braun V, Bühler A, Rottbauer W, et al. Hdac1-deficiency affects the cell cycle axis Cdc25-Cdk1 causing impaired G2/M phase progression and reduced cardiomyocyte proliferation in zebrafish. Biochem Biophys Res Commun [Internet]. 2023;665:98–106. Available from: 10.1016/j.bbrc.2023.04.116

46. Kobar K, Collett K, Prykhozhij S V., Berman JN. Zebrafish Cancer Predisposition Models. Front Cell Dev Biol. 2021;9(April):1–27.

47. Anastasaki C, Rauen KA, Patton EE. Continual low-level MEK inhibition ameliorates cardio-facio-cutaneous phenotypes in zebrafish. DMM Dis Model Mech. 2012;5(4):546–52.

48. Tucker B, Lardelli M. A rapid apoptosis assay measuring relative acridine orange fluorescence in zebrafish embryos. Zebrafish. 2007;4(2):113–6.

49. Greulich H, Erikson RL. An analysis of MEK1 signaling in cell proliferation and transformation. J Biol Chem [Internet]. 1998;273(21):13280–8. Available from: 10.1074/jbc.273.21.13280

50. García-Moreno JF, Lacerda R, da Costa PJ, Pereira M, Gama-Carvalho M, Matos P, et al. DIS3L2 knockdown impairs key oncogenic properties of colorectal cancer cells via the mTOR signaling pathway. Cell Mol Life Sci [Internet]. 2023;80(7):1–21. Available from: 10.1007/s00018-023-04833-5

51. Iyer S, Dhiman N, Zade SP, Mukherjee S, Singla N, Kumar M. Exposure to Tetrabutylammonium Bromide Impairs Cranial Neural Crest Specification, Neurogenic Program, and Brain Morphogenesis. ACS Chem Neurosci. 2023;14(10):1785–98.

52. Kimmel CB, Ballard WW, Kimmel SR, Ullmann B, Schilling TF. Stages of embryonic development of the zebrafish. Dev Dyn. 1995;203(3):253–310.

53. Schulte-Merker S, Ho RK, Herrmann BG, Nüsslein-Volhard C. The protein product of the zebrafish homologue of the mouse T gene is expressed in nuclei of the germ ring and the notochord of the early embryo. Development. 1992;116(4):1021–32.

54. Mahale S, Kumar M, Sharma A, Babu A, Ranjan S, Sachidanandan C, et al. The light intermediate chain 2 subpopulation of dynein regulates mitotic spindle orientation. Sci Rep [Internet]. 2016;6(1):1–16. Available from: 10.1038/s41598-016-0030-3

