## supplementary_tables for "DIS3L2 is essential for neural crest survival by modulating Akt signaling"

| **Gene** | **Forward primer Sequence** | **Reverse primer Sequence** | **Purpose** |
| --- | --- | --- | --- |
| *foxd3* | GCCTGGCAAAACTCCATTCG | TTGATAATCGACGCGGTGCT | ISH |
| *crestin* | ACCTGGAAATGCGACCCAAT | GCACTCTTCTCTGAAGCCGT | ISH |
| *twist1a* | GCCCGGTACATTGACTTCCT | TCAGGCCGAGAATCATGC | ISH |
| *sox10* | CACCACCCTCACGCTACAG | TCCACGTTACCGAAGTCGAG | ISH |
| *pax2a* | AGACCCCTACCTGACGTG | AGTCCAGGGTTCAGTGCT | ISH |
| *fgf8* | TTGCTACTATGCTCAGGTAACCA | GAGTAGCGGGTGCGTTTAGT | ISH |
| *bcl2a* | GAACTGGGGGCGGATCATT | ACGAAGGCATCCCAACCTC | qPCR |
| *bim* | TGACACGTCCAGAGAGCAA | CGGTGACCTCGACTGGTTAT | qPCR |
| *puma* | CCGAACCATTGCCACTCAAA | CACTTCCTGTTCTGTTCCTGA | qPCR |
| *foxo3a* | CGATGGCCTCTCTGACAACC | CGATGGCCTCTCTGACAACC | qPCR |
| *p21* | TGCAGAAGCTCAAAACATATTGTC | CTCAGACGCAAAGTCGAAGC | qPCR |
| *p27a* | AAAAGTGCGCGTCTCCAATG | CCGTACATCCTTGGCGAACT | qPCR |
| *p27b* | CGGGAATCACGACTGTAGGG | GGGTGTCGGACTCAATGGTT | qPCR |

**Fig. S1**

List of primers used in the study.
