## Supplementary figures and images for "DIS3L2 is essential for neural crest survival by modulating Akt signaling"

### supplementary_westernblots

**Figure 1E**

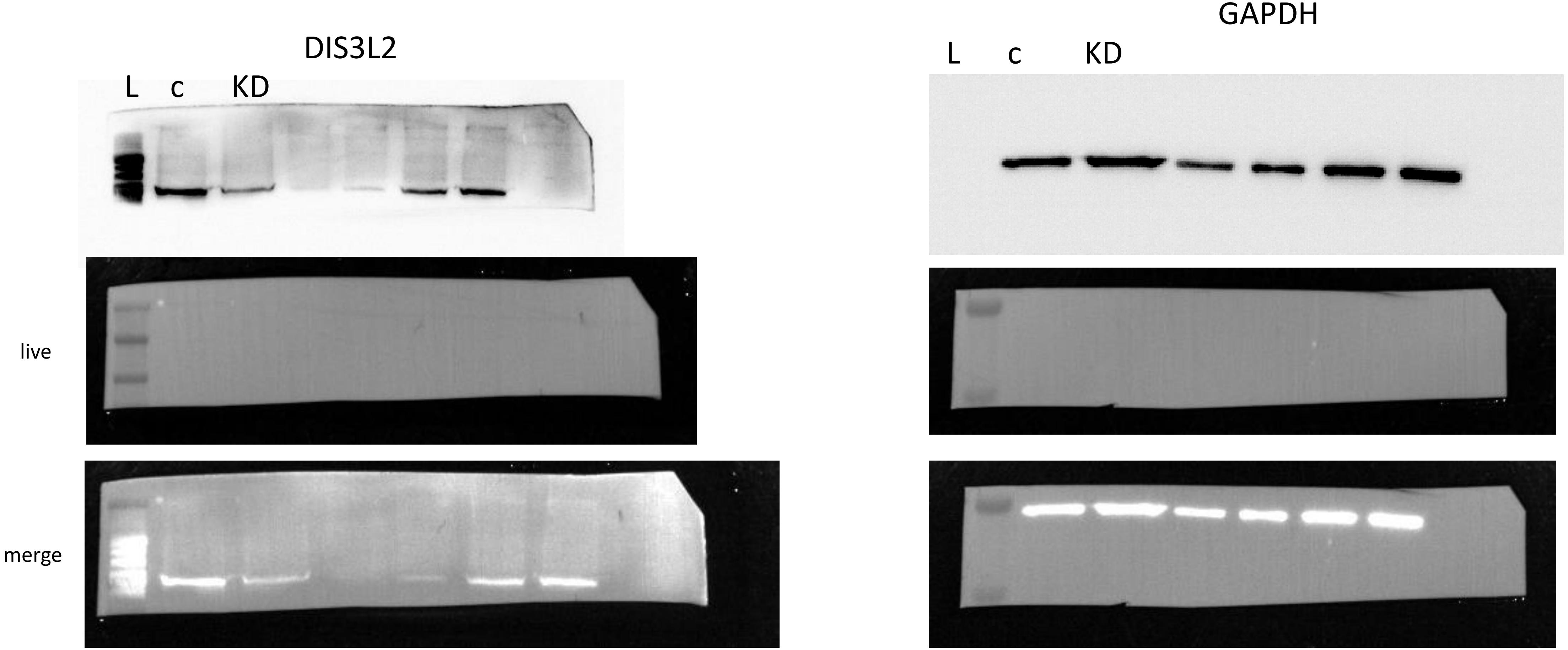

**Figure 3D**

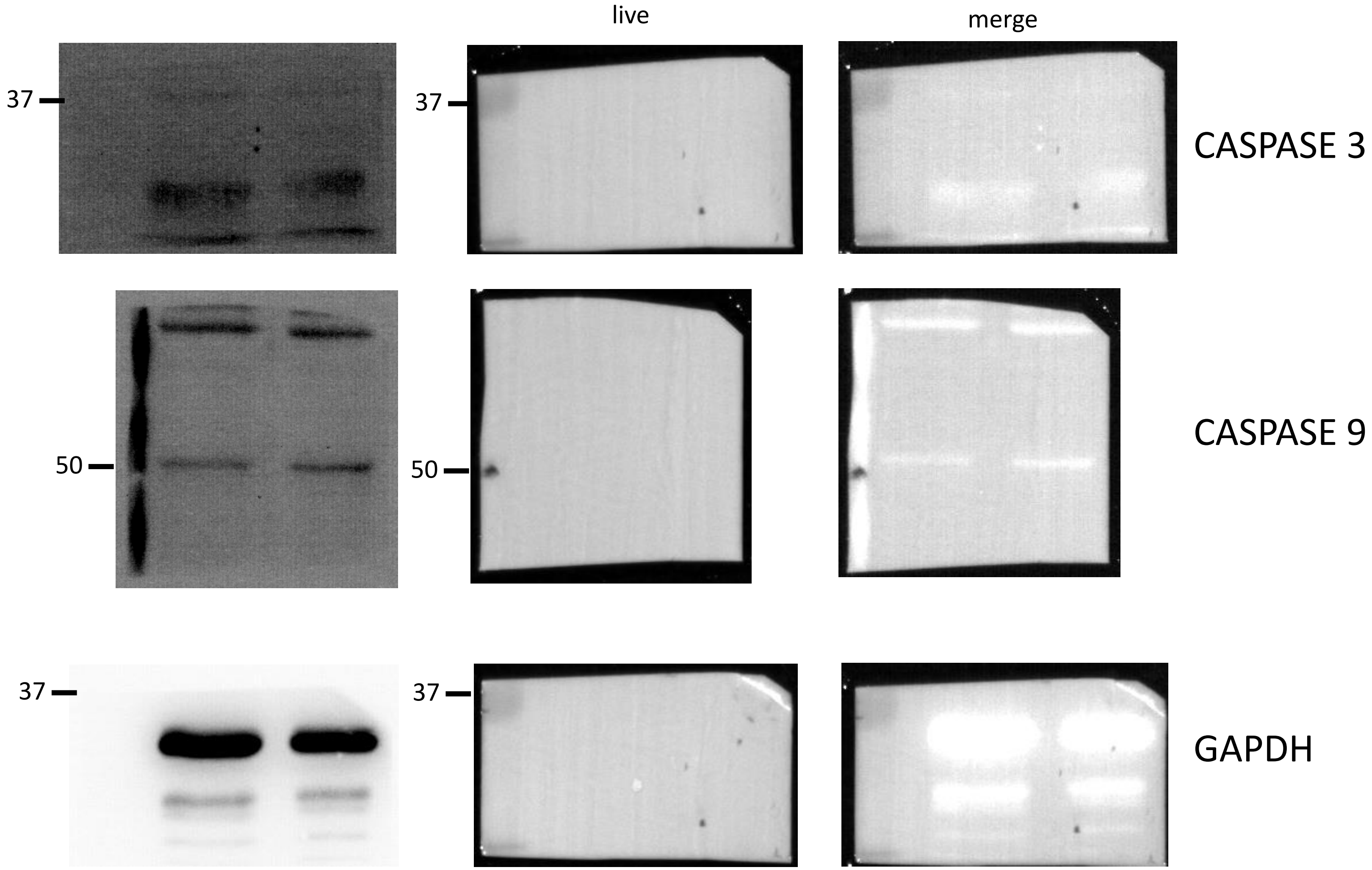

**Figure 3E**

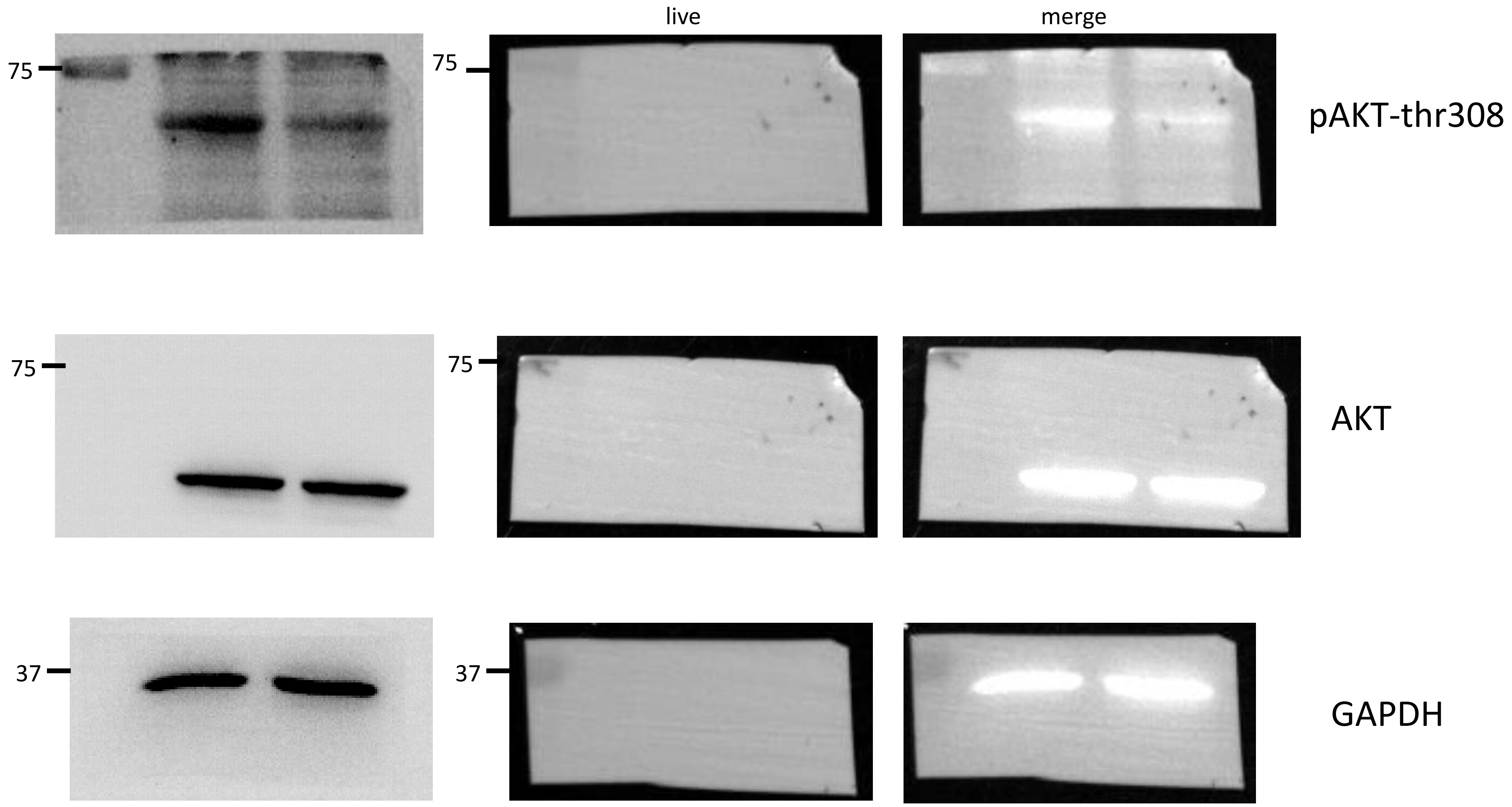

**Figure 3E**

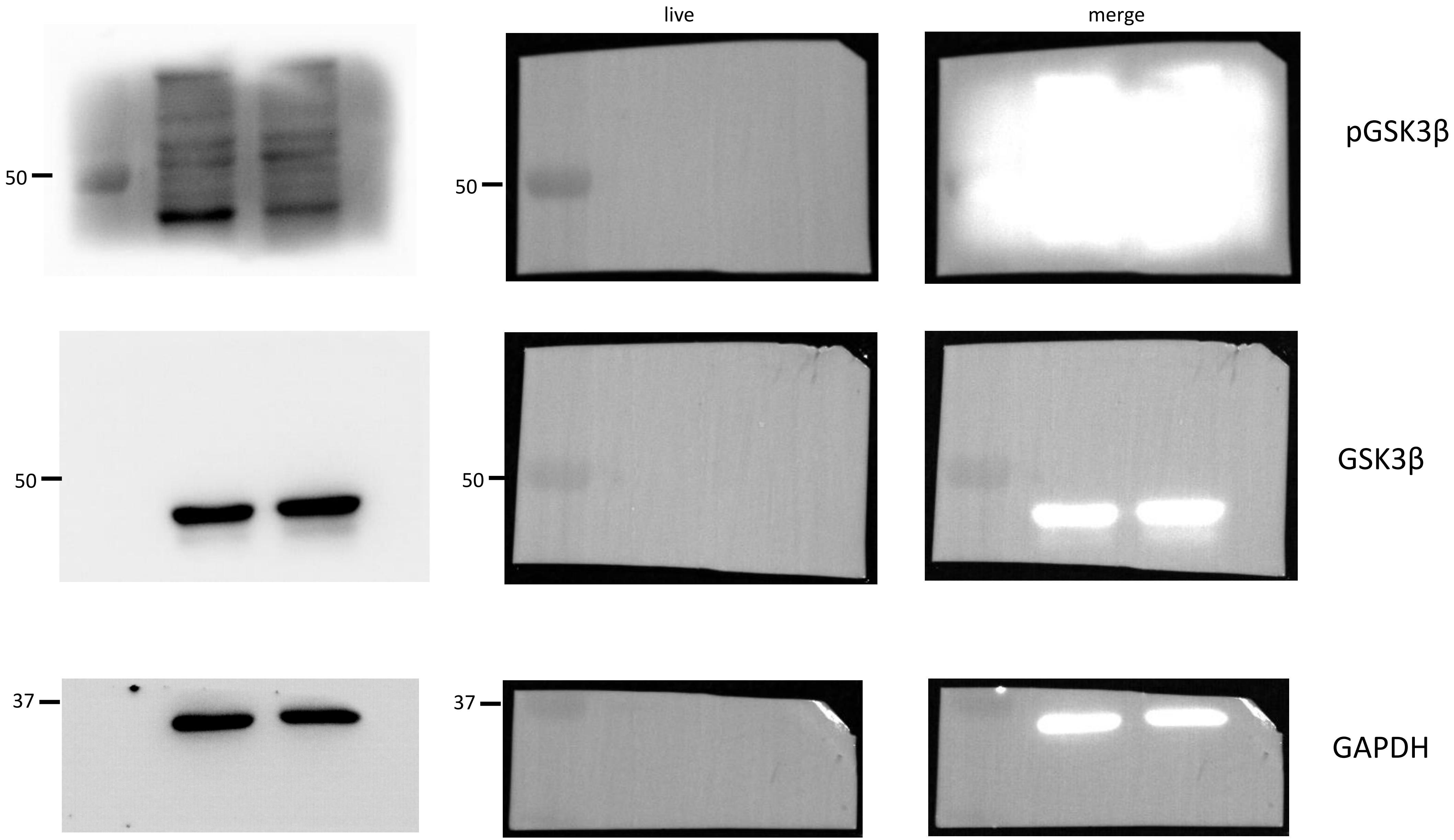

#### Figure 4E

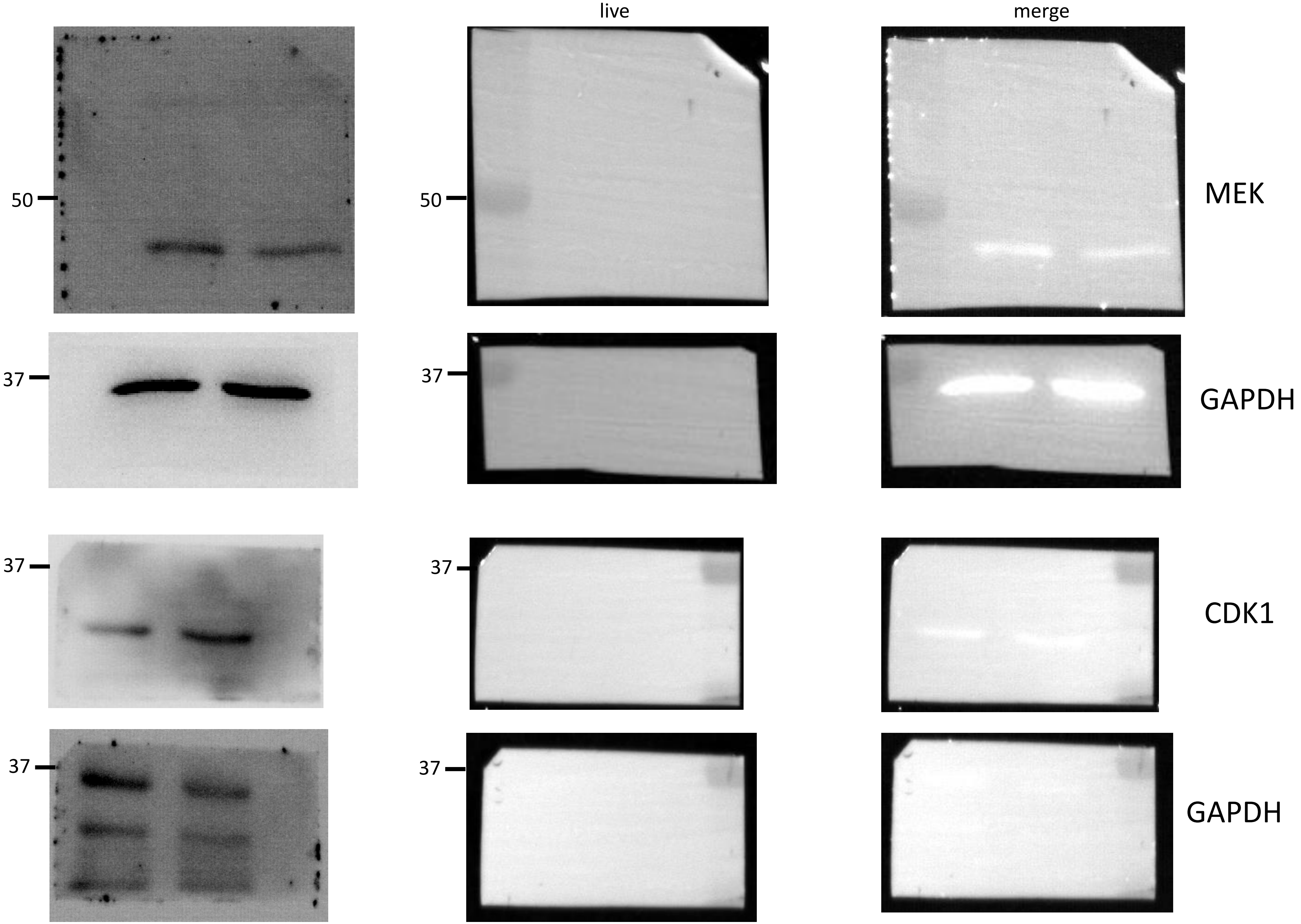
